# Nanosecond pulsed electric field ablates rabbit VX2 liver tumors in a non-thermal manner versus radiofrequency ablation

**DOI:** 10.1101/2022.08.16.504177

**Authors:** Qing-Gang Li, Zhen-Guo Liu, Ying Sun, Ya-Wen Zou, Xiao-Long Chen, Bin Wu, Xin-Hua Chen, Zhi-Gang Ren

**Affiliations:** Department of Infectious Diseases, the First Affiliated Hospital of Zhengzhou University, Zhengzhou 450052, China; Gene Hospital of Henan Province, Precision Medicine Center, the First Affiliated Hospital of Zhengzhou University, Zhengzhou 450052, China; Key Laboratory of Pulsed Power Translational Medicine of Zhejiang Province, Hangzhou 310003, China; Department of Hepatobiliary and Pancreatic Surgery, the First Affiliated Hospital, Zhejiang University School of Medicine, Hangzhou 310003, China

**Keywords:** Nanosecond pulsed electric fields, Ablation, Liver tumor

## Abstract

**Background:** Liver tumor remains an important cause of cancer-related death. Nanosecond pulsed electric fields (nsPEFs) are advantageous in the treatment of melanoma and pancreatic cancer, but their therapeutic application on liver tumors need to be further studied.

**Methods:** Hep3B cells were treated with nsPEFs. The biological behaviors of cells were detected by Cell Counting Kit-8, 5-ethynyl-20-deoxyuridine, and transmission electron microscopy (TEM) assays. *In vivo*, rabbit VX2 liver tumor models were ablated by ultrasound-guided nsPEFs and radiofrequency ablation (RFA). Contrast-enhanced ultrasound (CEUS) was used to evaluate the ablation effect. HE staining and Masson staining were used to evaluate the tissue morphology after ablation. Immunohistochemistry was performed to determine the expression of Ki67, proliferating cell nuclear antigen, and α-smooth muscle actin at different time points after ablation.

**Results:** The cell viability of Hep3B cells was continuously lower than that of the control group within 3 days after pulse treatment. The proliferation of Hep3B cells was significantly affected by nsPEFs. TEM showed that Hep3B cells underwent significant morphological changes after pulse treatment. *In vivo*, CEUS imaging showed that nsPEFs could completely ablate model rabbit VX2 liver tumors. After nsPEFs ablation, the area of tumor fibrosis and the expression of Ki67, proliferating cell nuclear antigen, and α-smooth muscle actin were decreased. However, after RFA, rabbit VX2 liver tumor tissue showed complete necrosis, but the expression of PCNA and α-smooth muscle actin did not decrease compared to the tumor group.

**Conclusions:** nsPEFs can induce Hep3B cells apoptosis and ablate rabbit VX2 liver tumors in a non-thermal manner versus RFA. The ultrasound contrast agent can monitor immediate effect of nsPEF ablation. This study provides a basis for the clinical study of nsPEFs ablation of liver cancer.

## Introduction

As population age [1] and risk factor changes [2], the number of new cases of primary liver cancer continues to increase. In 2018, the number of new cases of liver cancer was 841080 worldwide, accounting for 4.7% of all cancers, making liver cancer the sixth most common cancer type [3]. The number of deaths was 781631, accounting for 8.2% of all cancer deaths (fourth among all cancers) [3]. Although the global burden of hepatocellular carcinoma (HCC) could in theory be mitigated by universal HBV vaccination, the avoidance of environmental and lifestyle risk factors, and HCC surveillance in high-risk patients [4], the reality is not encouraging. In 2015, 257 million unvaccinated individuals, mostly in Asia and sub-Saharan Africa, were infected with HBV [5]. Therefore, the development of an effective treatment is still very important.

HCC patients suitable for resection still face a high risk of recurrence (up to 70% in 5 years) after surgery [6]. The number of HCC patients meeting the Milan criteria far exceeds the number of liver donors available for organ transplantation. For some patients with a single tumor less than 2 cm in diameter, radiofrequency ablation (RFA) can achieve the same therapeutic effect as resection [7]. Additionally, when the tumor is larger than 2 cm in diameter or located near important structures, the effect of RFA decreases [8]. Transarterial therapy and systemic therapy provide limited treatment for patients with intermediate and advanced disease.

Nanosecond pulsed electric fields (nsPEFs) can induce cell apoptosis by affecting the plasma membrane and intracellular structure [9,10], inhibiting tumor angiogenesis [11], reducing tumor metastasis [12], and producing antitumor immunity [13,14]. Quite different from irreversible electroporation which generally use parameters of microsecond pulse duration and 1500-V voltage, nsPEFs use comress the pulse width to nanosecond pusle duration under 30000-V voltage. nsPEF provide a sharp pulse with high energy. Compared with microsecond and millisecond pulse electric field, nsPEF showed superior tumor ablation potential. nsPEF lead to cell membrane structure polarization with super nanopore cause dielectric breakdown in short time periord without jeol heating accumulation, produce membrane electroporation without muscle convultion nor cardiac arrhythmia, nsPEF can penetrate into the cell, cause such as cytoskeleton depolymerization, intracellular calcium ion release and mitochondrial membrane potential dissipation. Based on the above features, nsPEF can effectively ablate tumors attached to high-risk structures without damaging blood vessels or bile ducts. *In vitro* and *in vivo* experiments of nsPEFs on liver tumors have made some progress in understanding the mechanism by which this treatment induces apoptosis, the immune response to treatment, and the effects of combination therapy [15].

To further verify the safety and effectiveness of nsPEF ablation for liver tumors, we conducted cell and animal experiments. This will provide a useful reference for the clinical application of nsPEFs in the multiple center clinical trials.

## Materials and methods

### Cell culture

The HCC cell line (Hep3B) was purchased from the Cell Bank of the Chinese Academy of Science (Shanghai, China). The cell line was cultured in Minimum Essential Medium (MEM Eagles with Earles Balanced Salts) supplemented with 10% fetal bovine serum (FBS), 100 U/mL penicillin and 100 mg/mL streptomycin. Cells were passaged three or four times per week after reaching confluence. The adherent cells were digested with 250 mg/mL trypsin for 1 min, and then the digested cells were transferred to a centrifuge tube. After centrifuge at 1000 r/min for 5 min, an appropriate amount of culture medium was added, and the cell suspension was mixed gently. For the pulse treated cells, we avoided centrifuge as much as possible.

### Induction of cell death by nsPEFs

A pulse generator device with two electrode needles (Fig. 1A) was provided by Hangzhou RuiDi Biological Technology Co., LTD (China). The field intensity was set at 10 kV/cm, the frequency was 2 Hz, the pulse number was set to 800, and the pulse duration was set at 0 and 300ns (the pulse duration was specified as the interval between the rising and falling edges at 90% amplitude). Waveforms were detected with a digital fluorescence oscilloscope. Hep3B cells were collected and re-suspended in fresh MEM containing 10% FBS, and the cell concentration was adjusted to 5.0 × 10^6^ cells/mL. A 500 μL aliquot of cell suspension was placed in a 0.1 mL colorimeter (Fig. 1B) and exposed to nsPEFs with pulse duration of 0 and 300 ns. The whole experiment was repeated three times.

**Fig. 1.**
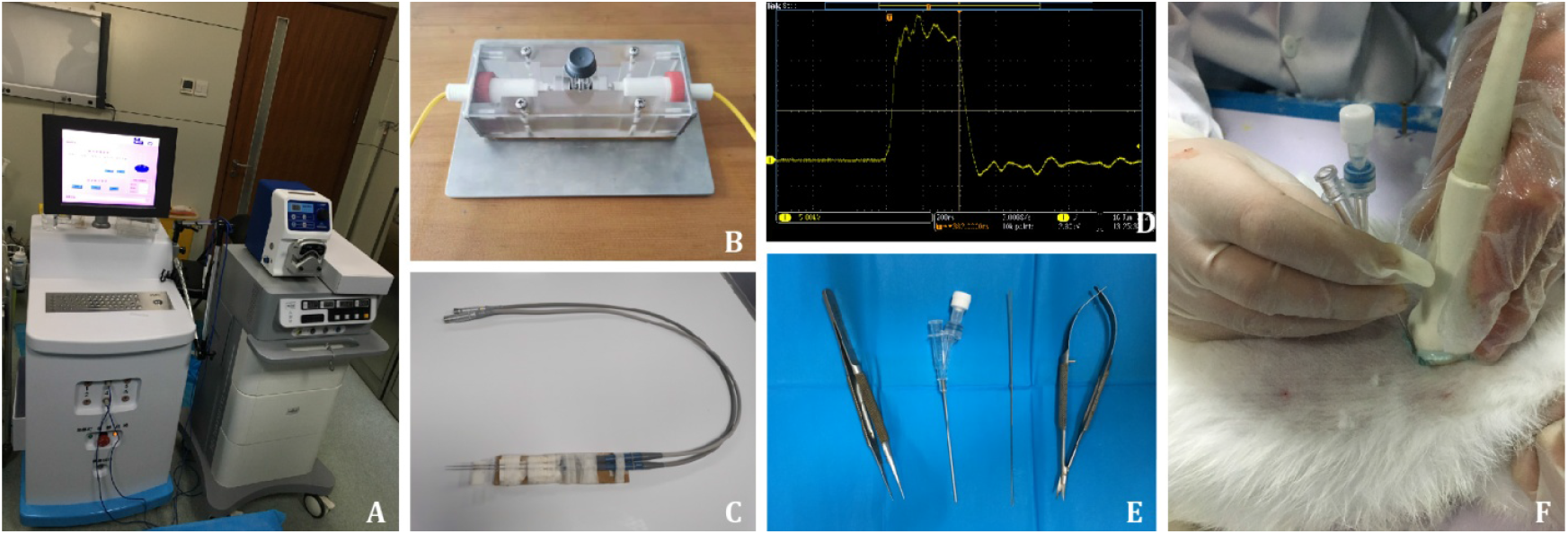
Ablation apparatus and the inoculation of rabbit VX2 liver tumors. **A:** The nsPEF ablation instrument is on the left, and the radiofrequency ablation instrument is on the right. **B:** Device for treating cells with nsPEFs. **C:** Ablation electrodes in animal experiments. **D:** The pulse waveform generated by nsPEFs. **E:** Preparation of the needles and tweezers. **F:** Percutaneous puncture under ultrasound guidance. nsPEFs: nanosecond pulsed electric fields.

### Cell viability

We evaluated the effect of nsPEFs on Hep3B cells with Cell Counting Kit-8 (CCK-8, Dojindo Laboratories, Shanghai, China). Hep3B cells treated with nsPEFs were collected and counted, and 100 μL of MEM containing 1 × 10^5^ Hep3B cells was finally added to the 96-well plate. At 24, 48, and 72 h after initiation of cell culture, 10 μL of CCK-8 reagent was added to the 96-well plate. After the plate was incubated at 37 °C in 5% CO_2_ for 1 h, the results were measured with a micro-plate reader (ELx800, BioTek Instruments, Winooski, VT, USA).

### EdU assay

A 5-ethynyl-20-deoxyuridine (EdU) assay kit (RiboBio, Guangzhou, China) was used to assess the cell proliferation ability. Hep3B cells were inoculated into confocal plates at a density of 10 × 10^5^ cells per well, followed by the addition of 50 μmol/L EdU buffer and incubation at 37 °C in 5% CO_2_ for 2 h. After fixing with 4% formaldehyde for 0.5 h, 0.1% Triton X-100 was added for 20 min to permeabilize the cells. Finally, EdU solution was added, and Hoechst was used to stain the nuclei.

### Observation of cell ultra-structure by transmission electron microscopy

The treated cells were immobilized with 2.5% phosphate buffered saline (PBS)-glutaraldehyde for 1-2 h and then washed with PBS to remove the fixative. The samples were soaked with PBS containing 1% OsO_4_ for 2 to 3 h and washed with PBS. Next, the samples were dehydrated with different grades of ethanol. After three washes with 100% acetone, the sample was embedded in Epon 812. Ultrathin slices (100 nm) were obtained using a Leica EM UC7 (Leica, Wetzlar, Germany). Finally, stained sections were observed using a Hitachi-7650 transmission electron microscope (HITACHI 7650, Tokyo, Japan).

### Animals and ethics statement

All animal experiments were in accordance with the guidelines for the management of animals for medical experiments issued by the Ministry of Health of the People’s Republic of China (No. 55) and approved by Experimental Animal Care and Ethics Committee of the First Affiliated Hospital of Zhengzhou University (KS-2017-KY-002). By anesthetizing animals with an appropriate amount of sodium pentobarbital, killing animals with an excess of sodium pentobarbital, and making every effort to minimize suffering.

### Ablation of normal rat liver tissue

After the experimental scheme was approved, animal care and feeding were carried out according to the ethical standards of the Animal Protection and Use Committee of Zhengzhou University. Six 8-week-old clean Sprague-Dawley rats (weighing 300-450 g) were purchased, and the rats were maintained in the Animal Experimental Center of Zhengzhou University. They were fed ad libitum under conditions of temperatures ranging between 22–24°C in a house with controlled 12-hour light and 12-hour dark cycle. After anesthesia, the rats were placed in the supine position and fixed on the operating table. After skin disinfection and sheet laying, an incision of approximately 1.5 cm was made along the midline of the abdomen of the lower sternal manubrium. Two electrodes (Fig. 1C) of the nsPEFs were inserted on both sides of the liver lobe with a spacing of 1 cm. The pulse parameters were set to a field intensity of 10 kV/cm, frequency of 2 Hz, pulse number of 800, and pulse duration of 300 ns (Fig. 1D). Twenty-four h after ablation, the fused tissue was removed for hematoxylin-eosin (HE) staining.

### Establishment of VX2 liver tumor model

A total of 42 healthy New Zealand white rabbits were purchased from the Zhengzhou University Animal Experiment Center. The lights were off without dim light phases from 7 p.m. to 7 a.m. The room temperature was 66°F (18°C) ± 4°F and the humidity was in the range of 40%-60%. VX2 tumor cells were injected into the hind limbs of anesthetized rabbits by intramuscular injection. When the tumor diameter reached 2 cm, the tumor tissue was removed from the rabbits with strict aseptic procedures and stored in liquid nitrogen. When tumor tissue was needed for experiments, it was thawed quickly at 36 °C and cut into cubes of 2 mm^3^. The necessary surgical instruments were prepared as shown in Fig. 1E. New Zealand white rabbits were anesthetized with 3% pentobarbital at a dosage of 30 mg/kg. After scraping the rabbit abdomen with povidone iodine solution (Fig. 1F), a small incision was made under the xiphoid process of the rabbit to expose the left medial lobe of the liver. VX2 tumor blocks were inserted into the liver parenchyma with a puncture needle under ultrasound guidance (Fig. 1F). A gelatin sponge was used to compress the liver surface to achieve hemostasis if needed. A daily dose of 50000 IU/kg sodium penicillin was administered for three days after inoculation. When ultrasound examination showed a tumor diameter of 1.5-2.0 cm, the model was considered to have been successfully established. After the rabbit was anesthetized, the sonographer inserted two electrodes around the tumor under ultrasound guidance. Six healthy rabbits were used as the normal control group. The successful inoculation of 30 rabbits was divided into five groups. The first four groups received nsPEF ablation and the final group received RFA. The parameters of nanosecond pulse were the same as those of rat ablation. The first four groups of rabbits were sacrificed at days 1, 2, 5, and 7 after ablation, and tumor tissue was collected. Five days after treatment, tumor tissue was taken from rabbits treated with RFA. In addition, six rabbits were successfully inoculated as the tumor group, i.e. electrodes were inserted but not electrified.

### Contrast-enhanced ultrasound (CEUS) evaluation

The ultrasound probe was placed at the level of the rabbit’s accessory xiphoid process. Two-dimensional ultrasound was used to determine the location and echogenicity of the tumor. The microbubble contrast agent was injected via an auricular vein. Thirty min after the ablation, CEUS was used to observe whether there were any lesions with residual abnormal blood perfusion in the ablation area.

### Histological assays

The tumor tissue was fixed in 4% paraformaldehyde and placed on a shaker overnight. The fixed tissue was embedded in paraffin, cut into 4 μm sections and subjected to HE and Masson’s trichrome staining. The area ratio of nuclei to the entire field of vision was calculated. The percent area of fibrosis was measured in the Masson-stained sections.

### Immunohistochemistry

Cancer cell proliferation was quantified by immunohistochemical analysis of Ki67 and proliferating cell nuclear antigen (PCNA). In addition, the activation of hepatic stellate cells was evaluated by immunohistochemical analysis of alpha-smooth muscle actin (α-SMA). All sections were reviewed by two pathologists, and each section was semiquantified on the basis of five different fields at high magnification.

### Statistical analysis

GraphPad Prism 8 software (La Jolla, CA, USA) was used to perform the statistical analysis. Continuous data were reported as mean ± standard deviation (SD). One-way analysis of variance (ANOVA) was used for comparisons of three or more groups. *P* < 0.05 was considered statistically significant.

## Results

### nsPEFs significantly inhibited the viability and proliferation of Hep3B cells

For Hep3B cells, cell viability was significantly decreased at 24, 48, and 72 h after nsPEF treatment (Fig. 2A). Compared with the control group, the cell viability of Hep3B cells at 24 h was 0.2494±0.0342 (*P* < 0.01), the cell viability of Hep3B cells at 48 h was 0.30534±0.0427 (*P* < 0.01), and the cell viability of Hep3B cells at 72 h was 0.4732±0.1150 (*P* < 0.01).

**Fig. 2.**
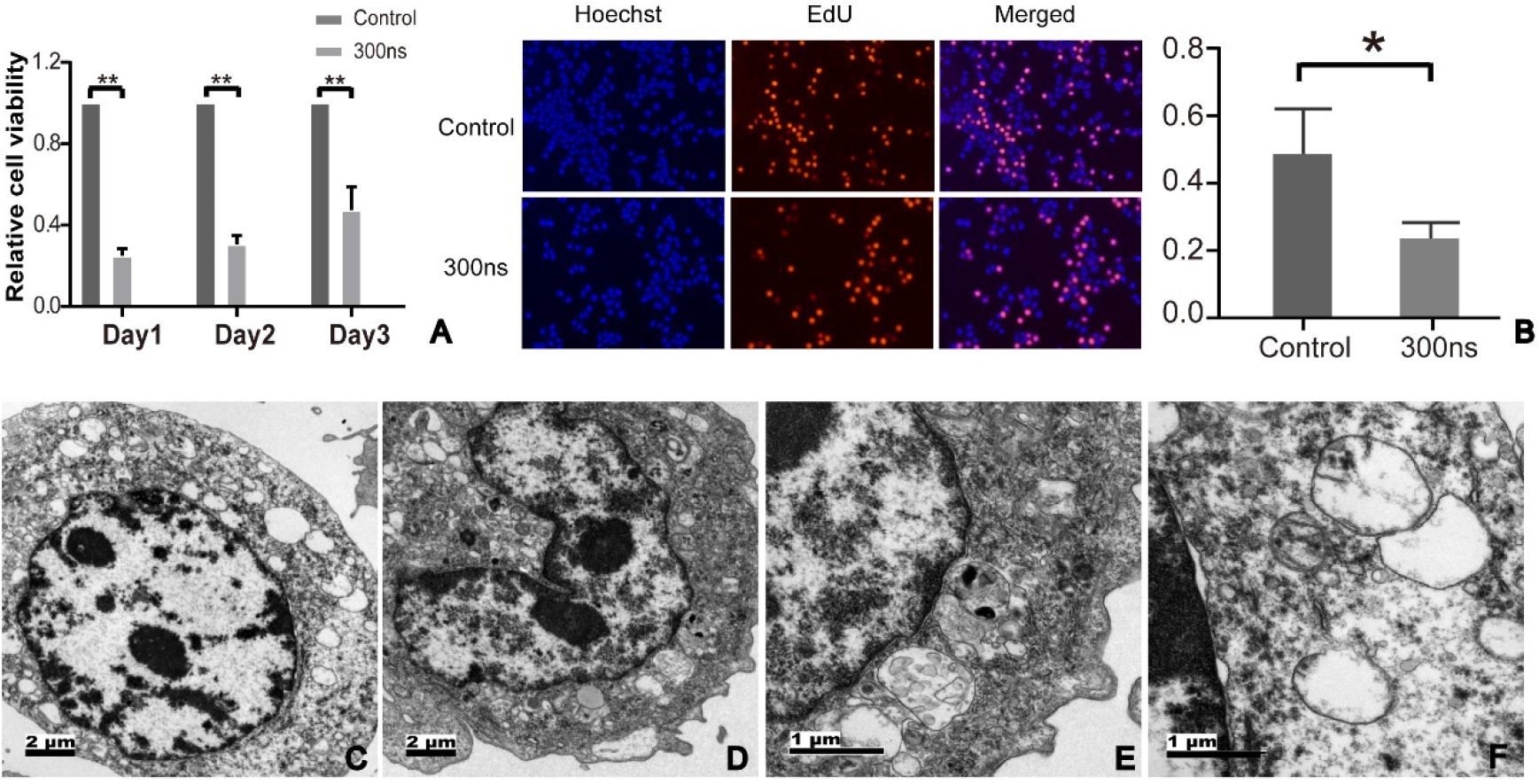
Hep3B cell behavior experiments. **A:** Hep3B cells were treated in cuvettes as shown in Fig. 1B with 2.5 × 10^6^ cells with pulses with durations of 0 and 300 ns and pulse numbers of 800, respectively. At days 1, 2, and 3 after treatment cell viability was determined by the CCK-8 assay. **B:** Hep3B cells were treated with pulsed electric field under the above parameters and inoculated into the confocal plate at a density of 10 × 10^5^ cells per well, and then a cell proliferation experiment was conducted. **C-F:** Hep3B cells treated with the pulsed electric field for 2 h were observed by transmission electron microscopy (**C** and **D**, original magnification × 2500; **E** and **F**, original magnification × 10000). Data are expressed as the mean ± SD. * *P* < 0.05; ** *P* < 0.01; *** *P* < 0.001. nsPEFs: nanosecond pulsed electric fields.

As shown in Fig. 2B, nsPEFs showed a significant inhibitory effect on the proliferation of Hep3B cells. Compared with the control group (0.4860±0.1346), the proliferation capacity of Hep3B cells treated with pulse was only 0.2369±0.0470 (*P* < 0.05).

### nsPEFs significantly affected the morphology of Hep3B cells

The morphology of tumor cells treated with nsPEFs was observed by transmission electron microscopy. Fig. 2C and D showed nuclear shrinkage, chromatin condensation, and margination. Fig. 2E and F showed enlarged, proliferated, and vacuolated mitochondria as well as expansion of the endoplasmic reticulum lumen.

### nsPEFs can preserve the approximate structure of the normal hepatic lobule

In normal rats, after nsPEF ablation, most hepatocytes were large and swollen, with pale eosinophilic cytoplasm, nucleolysis and necrosis (Fig. 3). At the same time, blood cells were seen at the boundary of the ablation zone. It should be noted that the portal vein and bile duct in the ablation zone were intact, and the general structure of the hepatic lobule was present.

**Fig. 3.**
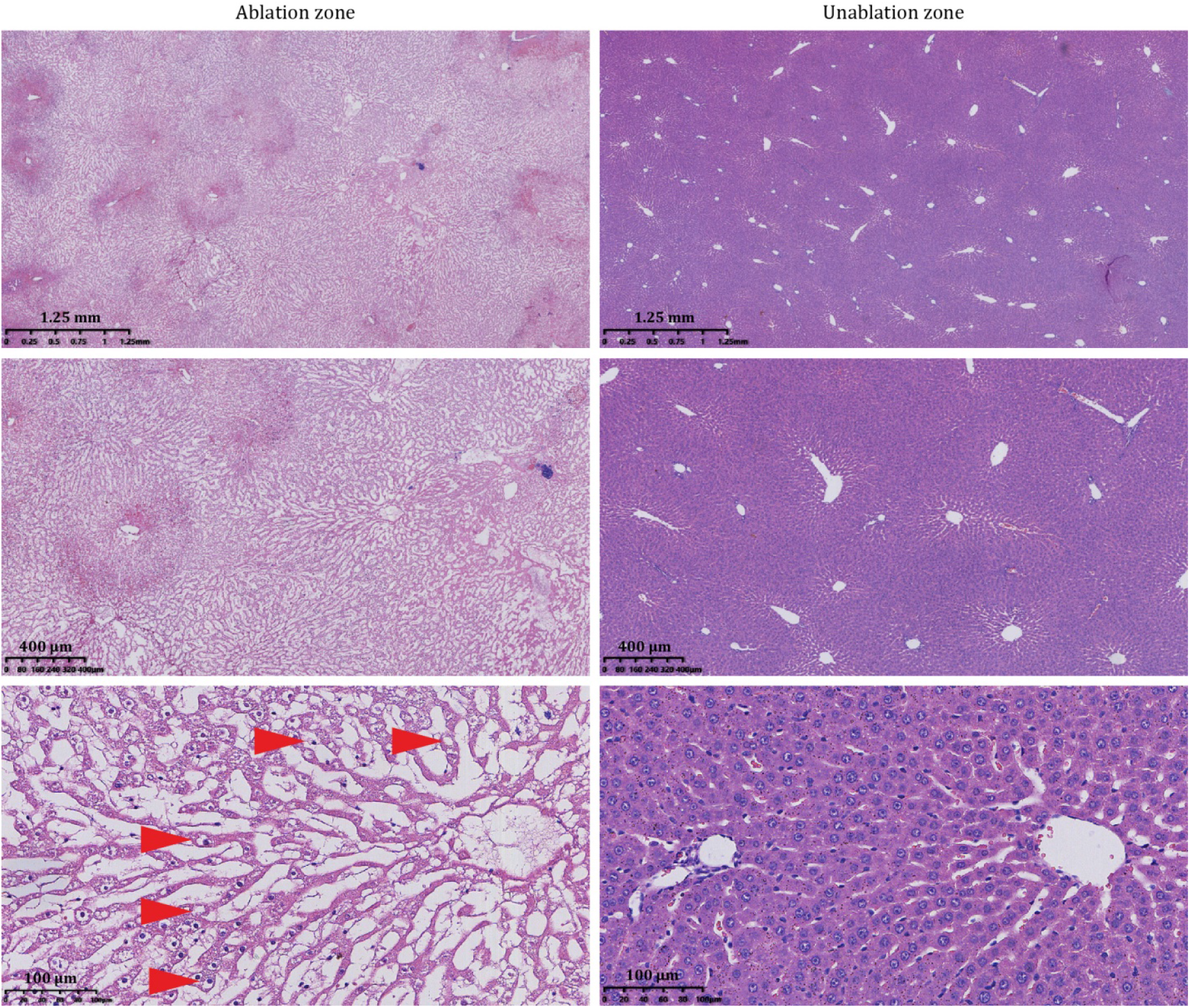
HE-stained sections of nsPEF-ablated and unablated regions of normal rat livers(n=6). The two electrodes of nsPEFs were inserted into the two sides of the liver lobe of anesthetized healthy rats at a distance of 1 cm. Pulse parameters were set as field intensity of 10 kV/cm, frequency of 2 Hz, pulse number of 800 times, pulse duration of 300 ns. At 24 h after ablation, HE staining was performed on the melted tissue. The light-colored areas in the image on the left are ablated areas, and the unablated areas are in the image on the right. As the red arrowheads show, at 24 h after nsPEF ablation, most hepatocytes were large and swollen, with pale eosinophilic cytoplasm, nucleolysis and necrosis. nsPEFs: nanosecond pulsed electric fields.

### The rabbit VX2 liver tumor was completely ablated by nsPEFs

The effectiveness and safety of nsPEFs were verified by animal experiments *in vivo*. The field strength was set to 10 kV/cm, and the pulse duration was set to 300 ns, the pulse number was set to 800. Fig. 4A-D showed representative ultrasonic images taken before and after nsPEF ablation. The CEUS image (Fig. 4A and B) of the tumor before nsPEF ablation presents a typical “fast-in and fast-out” appearance. The two-needle electrodes can be identified during ablation (Fig. 4C). After ablation, CEUS showed that there were no fast-in and fast-out enhanced lesions in the ablation area, indicating that a good ablation effect was achieved in the early stage (Fig. 4D). Fig. 4E and F represent CEUS images before and after RFA. In Fig. 4E, the heterogeneous enhanced lesion can be seen, while in Fig. 4F, no enhanced lesion can be seen. Obviously, CEUS results showed that the nsPEFs achieved a complete ablation effect.

**Fig. 4.**
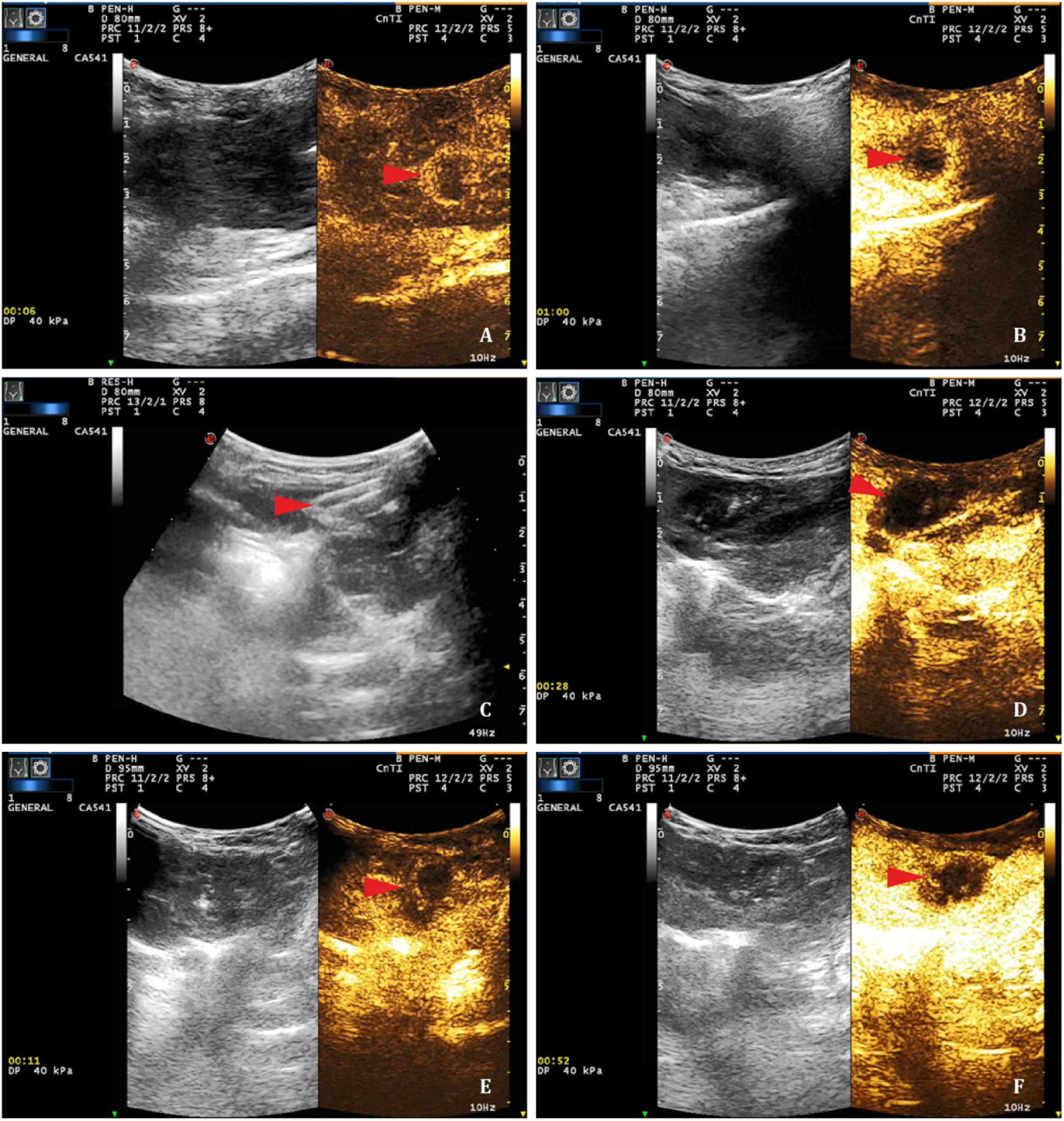
Ultrasound images of nsPEFs and RFA of VX2 liver tumor models. After the rabbit was anesthetized, the sonographer inserted two electrodes around the tumor under ultrasound guidance. Pulse parameters were set as field intensity of 10 kV/cm, frequency of 2 Hz, pulse number of 800 times, pulse duration of 300 ns. The ultrasonic probe was placed at the level of the xiphoid process. After 30 min ablation, contrast-enhanced ultrasound was performed to observe whether there was abnormal blood perfusion in the ablation area (red arrowheads). **A** and **B** represent ultrasound images taken prior to ablation. **C** shows the two electrodes of the nanosecond pulse ablation system. **D** represents the ultrasound image 30 min after nsPEF ablation. **E** and **F** represent the ultrasound images before and 30 min after RFA, respectively. nsPEFs: nanosecond pulsed electric fields; RFA: radiofrequency ablation.

### The expression of Ki67, PCNA, and α-SMA in liver tumors was significantly decreased by nsPEFs

As shown in Fig. 5A, on HE staining, the untreated tumor cells were densely packed with purplish-blue nuclei and pale purple cytoplasm. At days 1 (0.8490±0.0523, *P* < 0.01) and 2 (0.6959±0.0411, *P* < 0.001) after pulsed treatment, chromatin concentration and cytoplasm reduction were observed in some tumor cells. After treatment for 5 (0.6003±0.1112, *P* < 0.001) and 7 (0.4872±0.0468, *P* < 0.001) days, almost all tumor cells were extremely contracted and hyperchromatic, and the space between the cells became much larger. After RFA, liver tumor cells demonstrated marked reddish-purple staining resulting from nuclear fragmentation and cell necrosis (Fig. 5B).

**Fig. 5.**
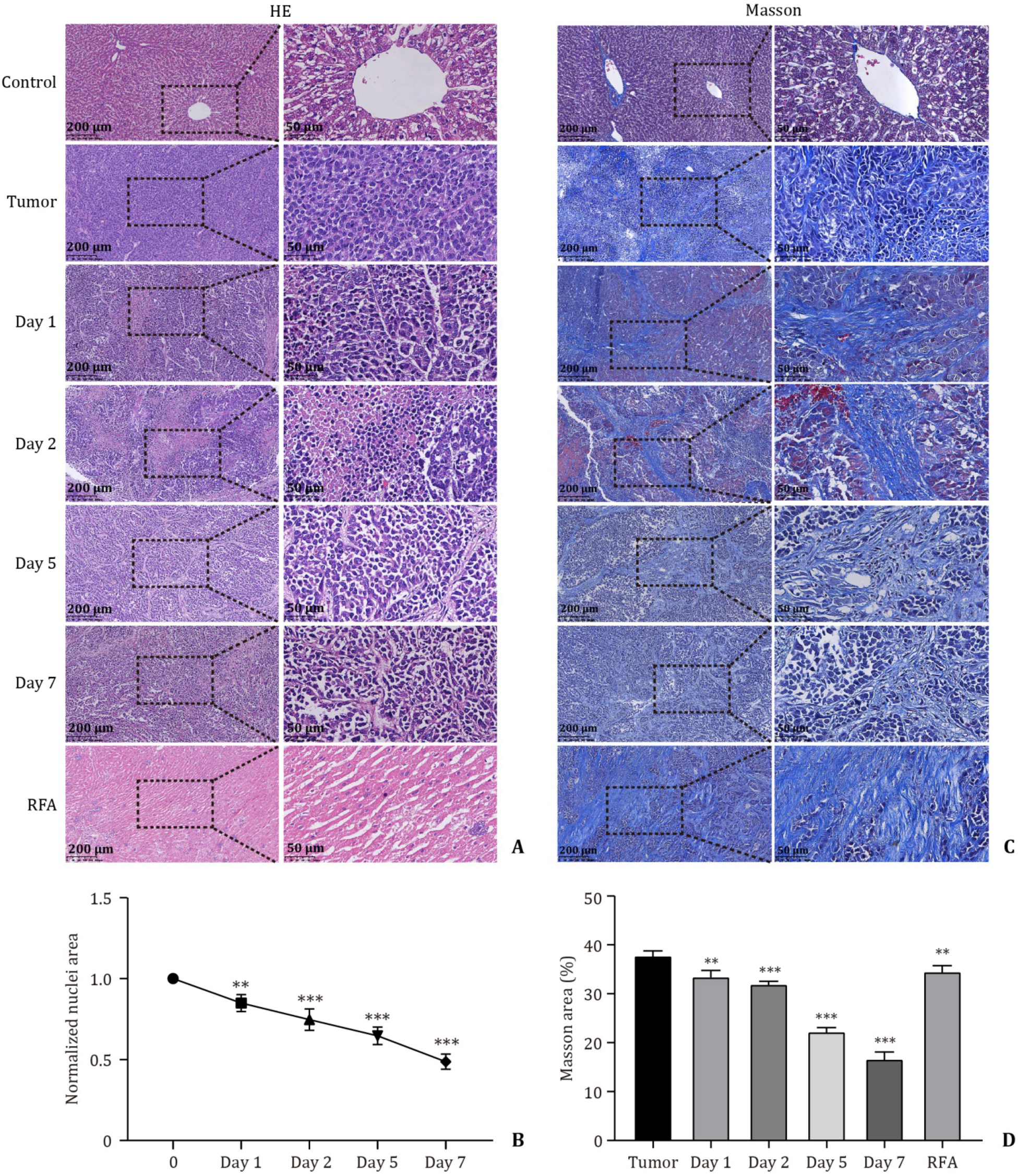
HE staining and Masson staining images and analysis of tumor tissue after ablation. **A** and **C:** HE and Masson staining were performed on the tissues at days 1, 2, 5 and 7 after nsPEF ablation and 5 days after RFA (*n* = 6). **B:** The percentage of nuclear area was calculated in the total high magnification field at different time points and groups (Days 1, 2, 5, and 7 were compared with the hypothesis at 0). **D:** The IOD percentage of the Masson staining area was calculated at different time points and groups (Days 1, 2, 5, and 7 and RFA groups were each compared with the tumor group). Data are expressed as the mean ± SD. ** *P* < 0.01; *** *P* < 0.001. nsPEFs: nanosecond pulsed electric fields; RFA: radiofrequency ablation; IOD: integrated optical density.

As shown in Fig. 5C, compared with that in the tumor group, the area of fibrosis in the tissues decreased in 1 (0.3356±0.0122, *P* < 0.01) and 2 (0.3204±0.0051, *P* < 0.001) days after nsPEF ablation and decreased significantly in 5 (0.2229±0.0079, P < 0.001) and 7 (0.1673±0.0135, *P* < 0.001) days. The RFA group (0.3358±0.0080, *P* < 0.01) also exhibited a reduction in fibrotic area, but not as large as that in the nsPEF group (Fig. 5D).

Compared with that in the tumor group (0.1591±0.0032; 0.0981±0.0031), the positive expression of Ki67 (Fig. 6) and PCNA (Fig. 6) in the nsPEF group was significantly decreased at days 5 (0.0580±0.0027, *P* < 0.001; 0.0477±0.0024, *P* < 0.01) and 7 (0.0478±0.0010, *P* < 0.001; 0.0229±0.0014, *P* < 0.01), and the positive expression of Ki67 in the RFA group was also decreased to some degree (0.0791±0.0058, *P* < 0.001). This indicated that the proliferation and expression of liver tumor tissues after nsPEF treatment were significantly inhibited. Similarly, α-SMA (Fig. 6), a myofibroblast marker associated with pathological liver stellate cell activation, was decreased significantly in 5 (0.4785±0.0021, *P* < 0.05) and 7 (0.0334±0.0005, *P* < 0.05) days after nsPEF treatment, while there was no statistically significant change in the RFA group (0.0862±0.0049, *P*> 0.05).

**Fig. 6.**
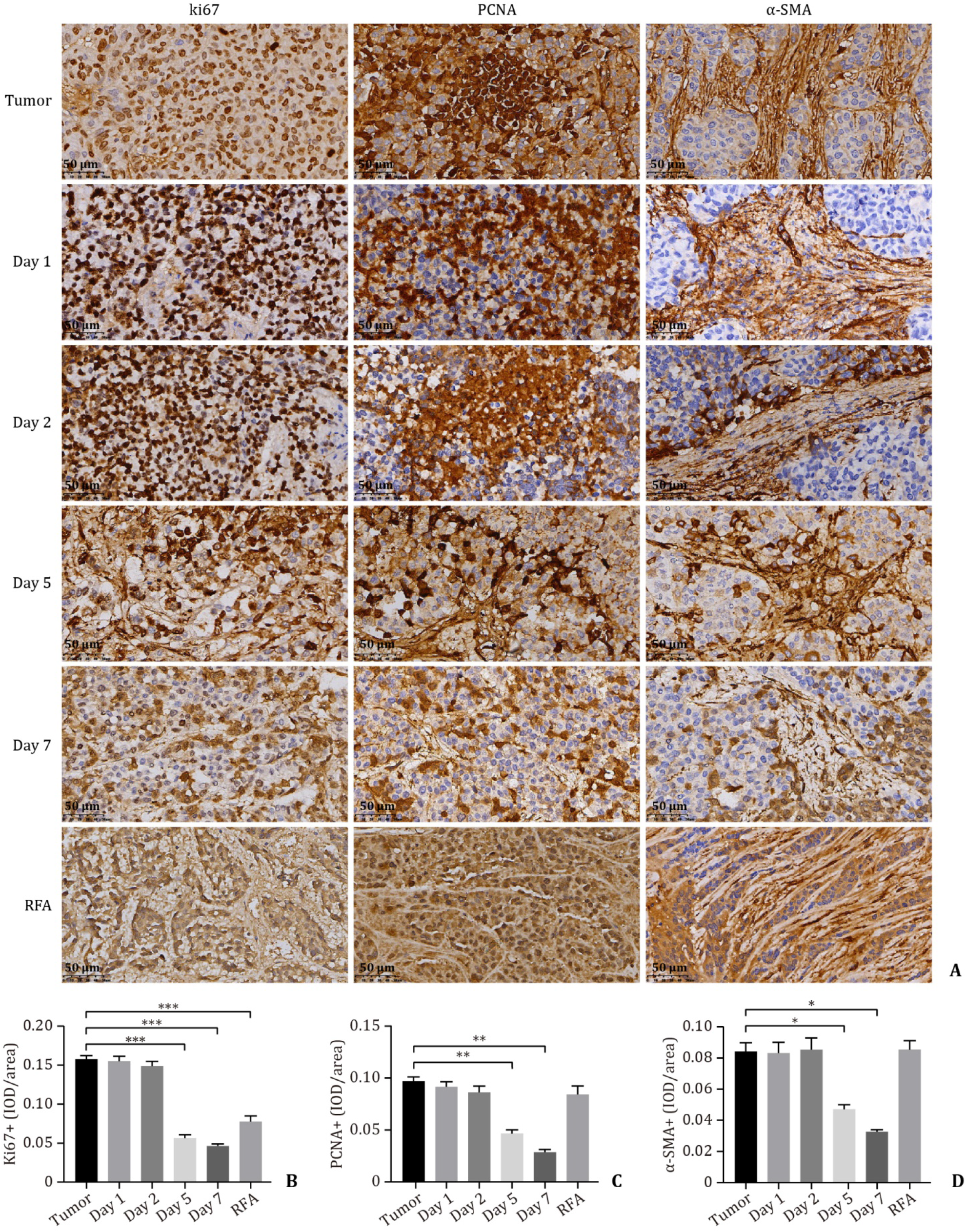
Immunohistochemistry of Ki67, PCNA, and α-SMA was performed on the VX2 tumor rabbit model at days 1, 2, 5 and 7 after nsPEF ablation. After 5 days RFA ablation, immunohistochemistry of Ki67, PCNA and α-SMA was performed on the rabbit VX2 liver tumor model. **A:** Representative immunohistochemical staining of Ki67, PCNA and α-SMA in liver sections from the different groups. **B-D:** Quantitative analysis of Ki67, PCNA and α-SMA staining in the different groups. Data are expressed as the mean ± SD. * *P* < 0.05; ** *P* < 0.01; *** *P* < 0.001. PCNA: proliferating cell nuclear antigen; α-SMA: alpha-smooth muscle actin; nsPEFs: nanosecond pulsed electric fields; RFA: radiofrequency ablation.

## Discussion

We not only investigated the effect of nanosecond pulses on Hep3B cells *in vitro* but also evaluated the ablation effect of nsPEFs in a rabbit VX2 liver tumor model. *In vitro* experiments showed that nsPEFs could affect the morphology of tumor cells and significantly inhibit the cell activity and proliferation of Hep3B cells. In the rabbit VX2 liver tumor model ablation experiment, we observed that the tumor could be completely ablated by nsPEFs. HE staining confirmed the occurrence of necrosis, and immunohistochemical analysis showed that the fibrotic area of the tumor was reduced, proliferation expression was decreased, and the activity of hepatic stellate cells was also decreased after ablation, which showed the unique advantages of nsPEFs. In experiments in normal rats, nsPEF ablation preserved the great vessels and bile ducts, a feature that can facilitate the repair of liver tissue.

The ablation effect of nsPEFs mainly depends on the field intensity, pulse duration, and pulse number. In *in vitro* studies [17,18], when two of the parameters remain unchanged, the changes in the other parameters lead to dose-dependent changes in the ablation effect. In addition, compared with bipolar nsPEFs, unipolar nsPEFs showed stronger membrane permeability changes and cell lethality [19]. Previous studies have proven the effect of high field intensity and high pulse number [18,20]. Considering the practical needs of clinical treatment, we adopted a low field intensity of 10 kV/cm. In addition, Chen et al. [21] also proved nsPEF achieve a complete ablation effect in liver tumor.

Under transmission electron microscopy, pulsed tumor cells exhibited signs of apoptosis: the chromatin in the nucleus was condensed, the micro-villi on the free surface of the cell membrane became thinner and reduced, the cytoplasm was condensed, the rough endoplasm reticulum expanded and swelled into vesicles, and the mitochondria became vacuolated. Previous studies [22,23] have found that nsPEFs with short pulse duration have a variety of effects on cells. Nanosecond pulses destroy the ultra-structure of the plasma membrane by affecting the charged molecules on its surface. In addition, rapid swelling of the mitochondria and rough endoplasm reticulum occurred in HCC in rats after nsPEF ablation [24]. nsPEFs can affect tumor viability by affecting the mitochondrial membrane potential [25,26], and plasma membrane permeability is also involved in apoptosis [27]. It should be noted that nsPEFs also damage micro-tubules [28], telomeres, cytoskeletal filaments, and other cellular structures [29,30]. It remains to be determined if the longer pulse duration have the same advantages as shorter duration that induce intracellular effects, which is superior to the microseoncd pulse which can only leak the membrane.

Ultrasound and CEUS are easy to operate and can display sensitively the blood perfusion of liver tumors in real-time. For example, liver tumors with a diameter of approximately 5 mm can be detected by CEUS [31]. Therefore, ultrasound examination effectively evaluates the inoperative and postoperative conditions of minimally invasive liver tumor ablation [32]. In the rabbit VX2 liver tumor model ablation experiment, postoperative ultrasound and CEUS showed that the ablated area was completely covered and that there was no residual active liver tumor. This suggests that ultrasound or CEUS can be used in clinical trials of nanosecond pulses to evaluate the tumor changes before and after tumor ablation. Chen et al. [20] ablated ectopic liver cancer mice with 900 pulses at 100 ns pulses and 68 kV/cm in a single treatment or in three treatment sessions, and the ultrasound results showed that 75% of primary HCC tumors had no recurrence within 9 months.

After nsPEF ablation, the tissue tended to necrotic within a few days, beside the ablation zone death, the treatment area showed also reduced fibrosis, proliferation, and expression and decreased production of active hepatic stellate cells. Therefore, nsPEFs may stimulate a protective response in organisms. Ren et al. [11] found that nsPEFs inhibited cell proliferation by inhibiting the NF-κB signaling pathway and down-regulated the expression of vascular endothelial growth factor and MMP family proteins by inhibiting the Wnt/β-catenin signaling pathway, thereby reducing tumor metastasis and invasion. In addition, the supporting structure of normal liver tissue remains intact after ablation [33]. Local intrahepatic pulse ablation can increase the level of hepatocyte growth factor (HGF), up-regulate platelet-derived growth factor in the ablation zone, activate the HGF/c-Met pathway, and promote whole-liver regeneration [34], all of which will promote liver tissue repair. This activity was confirmed by Chen et al. [13], who observed the emergence of new normal liver tissue 6 weeks after ablation. Notably, Nuccitelli et al. [35] demonstrated that nsPEF ablation can inhibit secondary tumor growth by triggering the production of CD8+ T-cells. Considering the effects of CD8+ T-cells, it will be valuable to determine if the longer duration exhibit intracellular effects and if these are critical for ablation and immunity as shown in rat HCC studies.

In conclusion, our study showed that nsPEFs could significantly affect the activity, proliferation, and cell morphology of Hep3B cells. Meanwhile, in the rabbit VX2 liver tumor model, compared with RFA, nsPEFs can induce cell apoptosis, reduce the area of tumor tissue fibrosis, and reduce tumor tissue proliferation expression. This study will provide ultrasound doctors with a minimally invasive med tumor ablation device. nsPEF does not produce Joule heat and can trigger activate immunity, it can be safely applied for tumors attached to large blood vessels. Therefore, it can break the forbidden zone of surgery and thermal ablation. nsPEF has advantages in enhanced recovery after surgery (ERAS) with more radical ablation effect and shorter hospital stay and less the economic burden.

